# HaploThread: A Scalable Integrated Desktop Platform for Constructing and Visualizing Haplotype Networks for Large-sample Sequences

**DOI:** 10.1101/2025.07.06.659816

**Authors:** Bo Xu, Lun Li, Cuiping Li, Anke Wang, Zhuojing Fan, Shuhui Song

## Abstract

This note announces HaploThread, a user-friendly GUI desktop software designed for haplotype network construction and visualization. HaploThread is written in C++ using the Qt library, integrating network visualization and multiple multi-threaded haplotype construction algorithms such as McAN and fastHaN (includes MSN, MJN and TCS) based on plugin mechanisms. It offers a straightforward approach to constructing and visualizing haplotype networks from large sample sets, and to extending functionality with plugins, facilitating the analysis of genetic variations and their evolutionary relationships. HaploThread is an open-source software released under the GNU General Public License (GPL). Its precompiled executables for Windows is freely available for download at https://ngdc.cncb.ac.cn/biocode/tool/BT007948.

## Introduction

Desktop software plays a crucial role in biological research by providing powerful computational tools with user-friendly graphical interfaces, enabling researchers to analyze complex biological data efficiently (Kumar and Dudley 2007). Favoured by biologists, it offers a balanced solution that combines ease of use, high performance, and data security—unlike command-line tools that require programming skills or web-based platforms that depend on internet access.

Particularly in population genetics, desktop software of haplotype network analysis is significant in inferring evolutionary relationships, tracking disease transmission, and studying population dynamics (Song, et al. 2020; Li, et al. 2024). However, traditional state-of-art desktop software for haplotype network construction and/or visualization, including TCS (Clement, et al. 2000), Network (https://www.fluxus-engineering.com/), PopART (Leigh and Bryant 2015), HapStar (Teacher and Griffiths 2011) and Hapsolutely (Vences, et al. 2024), only integrated single-threaded algorithms (Templeton, et al. 1992; Excoffier and Smouse 1994; Bandelt, et al. 1999; Matschiner 2016), which limits their efficiency when processing large datasets (**Table 1)**.

**Table 1.**
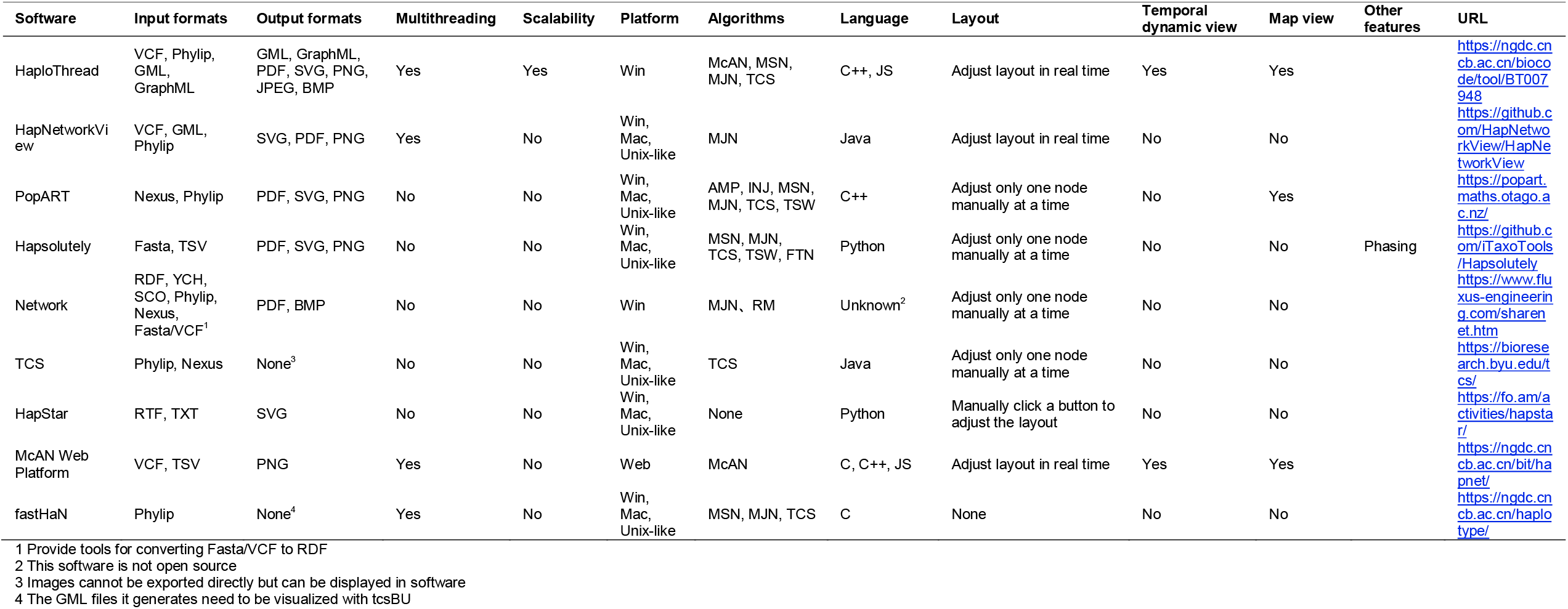
Comparison of programs for haplotype networks construction and/or visualization.

Currently known multi-threaded construction algorithms include McAN (Li, et al. 2023) and fastHaN (Chi, et al. 2023), which are faster than all existing algorithms even in single-threaded mode. Among them, McAN is currently the only program capable of processing sequences on the order of millions. Nevertheless, McAN provide only a simple command-line interface, which may appear daunting to beginning users, and an online web service, which depends on internet connection availability and stability and may bring concerns about privacy and security risks of user data.

To addresses these challenges, we developed HaploThread, a fully local software that integrate intuitive user interfaces with advanced multi-threaded algorithms, ensuring ease of use, data security, and computational efficiency in haplotype network construction and visualization. Built upon McAN, an algorithm from our previous work, and enhanced with the advanced features of fastHaN, HaploThread leverages the performance advantages of desktop environments and multi-threading to deliver rapid, scalable, and reliable analyses for evolutionary studies in population genetics.

### Functionality

HaploThread currently comprises two functional modules: the network construction module and the network visualization module (**Figure 1**). Each module is available in two distinct visual style of interface: a platform-dependent style and a web-based style.

**Figure 1.**
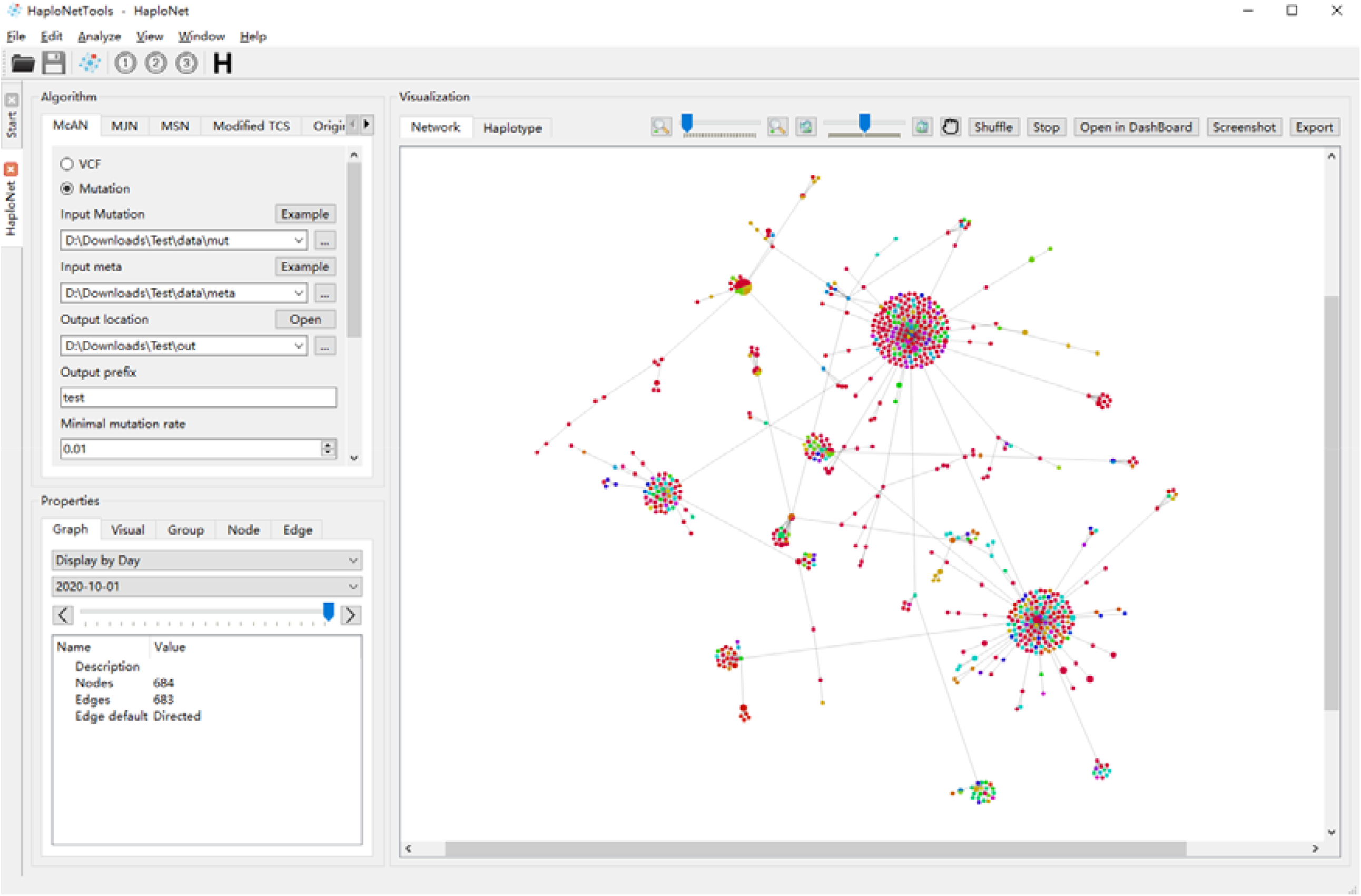
Screen shot for the interactive interface for network construction/visualization in HaploThread

The construction module processes sequence data in either VCF or PHYLIP format and generates the constructed network file in GraphML or GML format. Both styles of interfaces offer identical functionality, ensuring a consistent user experience. Users have the flexibility to choose a multi-threaded construction algorithm from McAN, TCS, MSN, or MJN, specify the required parameters, configure the number of computational threads based on available system resources, and start the process by clicking the “Run” button. The entire workflow is designed to be intuitive and user-friendly, requiring minimal explanation for efficient operation.

The visualization module is designed to open network files in GraphML or GML formats and employs a force-directed algorithm to layout the network. If users provide metadata, such as sampling date, geographic location, or clustering information, via a metadata file, all samples and network nodes are automatically assigned distinct colors according to the associated metadata, generating visualization color schemes that are available for user selection and application. Additionally, users can observe the temporal dynamics of the network by either automatically running or manually dragging the timeline.

The constructed network can be converted between GraphML and GML formats and exported as a PDF file or images in various formats, such as SVG, PNG, JPEG, or BMP.

The functionalities of the two styles of visualization differ slightly. The platform-dependent interface provides a visual browser for haplotype sequence differences, which can be displayed and selected in synchronization with the haplotype network. Users can customize the colour and font attributes of visual elements to suit their preferences. In contrast, the web-like interface offers a map browsing function, allowing nodes to be displayed on a world map when country-level metadata is available.

### Implementation

HaploThread software is written using the Qt cross-platform application framework (**Figure 2**). Compiled executables for Windows, macOS, and Linux are provided. It is architected with a plugin-based approach. Functional modules are designed in the form of plugins and can be integrated, updated or uninstalled independently of the core application, reflecting the flexibly and extensibility of the software.

**Figure 2.**
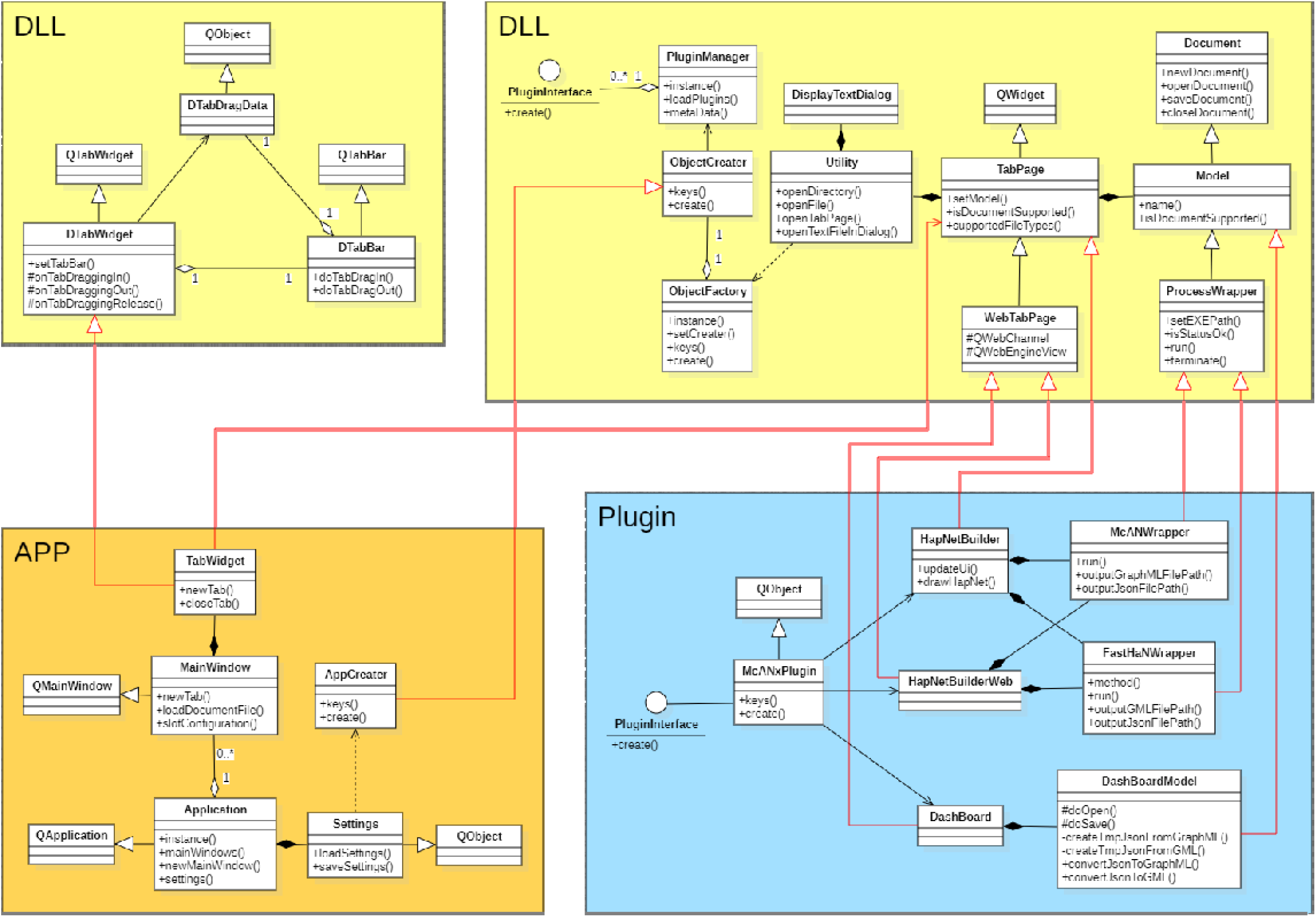
UML class diagram for the design of HaploThread.

The haplotype network construction module in HaploThread is implemented by integrating the software McAN and fastHaN into a plugin. The platform-dependent style of the module is developed using C++ and built on the Qt Widgets module. This approach ensures high performance and enhanced flexibility for desktop applications, and seamless consistency with the native styles of operating systems such as Windows, macOS, and Linux. The web-based style of the module is written using JavaScript and HTML, and it is embedded within the plugin via the Qt WebEngine module. This design provides users with an experience that is consistent with online web services.

Future improvements may involve the implementation of novel algorithms for network construction and layout, 3D visualization of networks, and expanding support for additional input and output formats.

### Performance Comparison

To evaluate the computational performance of different haplotype network construction tools, we conducted a benchmarking experiment using SARS-CoV-2 sequences downloaded from the RCoV19 database. Three datasets were prepared, containing 500, 1,000, and 5,000 sequences, respectively (**Data S1**). We tested five representative state-of-the-art desktop software tools: PopART, HaploThread, Hapsolutely, Network, and HapNetworkView. For a fair comparison, all tools were run in single-threaded mode regardless of their support for parallel computing.

To reflect typical usage scenarios for desktop applications, all tests were conducted on a consumer-grade laptop running Windows 10 Home Edition, equipped with an Intel Core i7-10510U CPU (1.80 GHz, 4 cores, 8 logical processors), 16 GB RAM, and a 512 GB SSD.

The results demonstrate that HaploThread consistently outperformed all other tools in terms of execution time across all three datasets (**Table S1**). For tools that offer multiple algorithms, only the fastest configuration was included in the comparison. Notably, on the 5,000-sequence dataset, HaploThread completed both network construction and visualization in just 23 seconds, while PopART, Hapsolutely, and Network were unable to finish the task within one hour. This highlights the superior efficiency and scalability of HaploThread in handling large-scale haplotype data under real-world computing conditions.

## Discussions and Conclusion

The rapid emergence and evolution of novel and outbreak-associated pathogens pose significant challenges to public health surveillance and epidemic control. Timely and accurate construction of haplotype networks is essential for tracking pathogen evolution, understanding transmission dynamics, and informing intervention strategies. Traditional haplotype network construction GUI desktop software often struggles to handle large-scale genomic datasets efficiently, limiting their applicability in urgent outbreak scenarios. HaploThread was specifically designed to address that problem.

Currently, a state-of-the-art desktop software for haplotype network construction and visualization, named HapNetworkView (Chi, et al. 2025), has been developed. It integrates a multithreaded MJN algorithm from fastHaN. In comparison, HaploThread offers broader advantages in terms of algorithmic diversity. In addition to supporting a multithreaded MJN algorithm, HaploThread integrates five state-of-the-art multithreaded haplotype network construction algorithms, including McAN, MJN, MSN, and two implementations of TCS. This comprehensive algorithm suite provides users with enhanced flexibility and computational efficiency, especially when analysing large-scale datasets, making HaploThread a versatile and powerful tool for haplotype network analysis. In addition to its algorithmic strengths, HaploThread also offers advanced visualization capabilities that enhance interpretability and user interaction. HaploThread not only enables colouring of nodes in the haplotype network based on geographic information, but also supports the dynamic display or concealment of nodes according to sampling time. These spatiotemporal visualization capabilities provide users with enhanced flexibility to explore evolutionary relationships and track the temporal and geographic spread of haplotypes, which is especially valuable in infectious disease research.

In summary, HaploThread simplifies the complex process of haplotype network analysis by integrating multiple methods of construction and visualization into a unified, user-friendly platform. Its intuitive design ensures accessibility for users with varying levels of expertise. Furthermore, the tool’s ability to handle large datasets, combined with its support for plugin mechanisms, facilitates community-driven improvements and customization, enhancing its adaptability to evolving research needs.

## Acknowledgements

This work was supported by the Key Collaborative Research Program of the Alliance of National and International·Science Organizations for the Belt·and·Road Regions (Grant No. ANSO-CR-KP-2022-09), and the National Natural Science Foundation of China (Grant No. 32270718, 32170678).

## Code Availability

Installation packages of HaploThread are freely available at https://ngdc.cncb.ac.cn/biocode/tool/BT007948 under the GNU General Public License (GPL).

## Data Availability

The datasets used for performance benchmarking in this study—comprising subsets of 500, 1,000, and 5,000 SARS-CoV-2 sequences from the RCoV19 database—are provided in **Supplementary Material 1**.

